# Avian ANP32B does not support influenza A virus polymerase and influenza A virus relies exclusively on ANP32A in chicken cells

**DOI:** 10.1101/512012

**Authors:** Jason S. Long, Alewo Idoko-Akoh, Bhakti Mistry, Daniel H. Goldhill, Ecco Staller, Jocelyn Schreyer, Craig Ross, Steve Goodbourn, Holly Shelton, Michael A. Skinner, Helen M. Sang, Mike J. McGrew, Wendy S. Barclay

**Affiliations:** Section of Molecular Virology, Imperial College London, St Mary’s Campus, London, W2 1PG, UK; The Roslin Institute and Royal (Dick) School of Veterinary Studies, University of Edinburgh, Easter Bush Campus, Midlothian, EH25 9RG, UK; Institute for Infection and Immunity, St. George’s, University of London, London, SW17 0RE, UK; The Pirbright Institute, Surrey, GU24 0NF, UK

**Keywords:** Influenza, polymerase, genome editing, disease, ANP32A, ANP32B, host pathogen interaction

## Abstract

Influenza A viruses (IAV) are subject to species barriers that prevent frequent zoonotic transmission and pandemics. One of these barriers is the poor activity of avian IAV polymerases in human cells. Differences between avian and mammalian ANP32 proteins underlie this host range barrier. Human ANP32A and ANP32B homologues both support function of human-adapted influenza polymerase but do not support efficient activity of avian IAV polymerase which requires avian ANP32A. We show here that avian ANP32B is evolutionarily distinct from mammalian ANP32B, and that chicken ANP32B does not support IAV polymerase activity even of human-adapted viruses. Consequently, IAV does not replicate in chicken cells that lack ANP32A. Amino acid differences in LRR5 domain accounted for the inactivity of chicken ANP32B. Transfer of these residues to chicken ANP32A abolished support of IAV polymerase. Understanding ANP32 function will help develop antiviral strategies and aid the design of influenza virus resistant genome edited chickens.

## Introduction

Influenza A viruses (IAV) infect a wide range of host species but originate from wild birds. Zoonotic transmission from the avian reservoir is initially restricted by host specific species barriers. Infection of new host species requires the virus to bind to cell surface receptors, utilise foreign host cellular proteins whilst evading host restriction factors in order to replicate its genome, and finally transmit between individuals of the new host.

The negative sense RNA genome of influenza A virus (IAV) is replicated in the cell nucleus using a virally encoded RNA-dependent RNA polymerase, a heterotrimer composed of the polymerase basic 1 (PB1), polymerase basic 2 (PB2) and polymerase acidic (PA) proteins together with nucleoprotein (NP) that surrounds the viral RNA, forming the viral ribonucleoprotein complex (vRNP)^1^.

Crucially, the viral polymerase must co-opt host factors to carry out transcription and replication^1^. The PB2 subunit is a major determinant of the host restriction of the viral polymerase^2^. Avian IAV polymerases typically contain a glutamic acid at position 627 of PB2, and mutation to a lysine, the typical residue at this position in mammalian-adapted PB2^3^, can adapt the avian polymerase to function efficiently in mammalian cells. We have suggested that the restriction of avian IAV polymerase is due to a species specific difference in host protein ANP32A^4^. Avian ANP32A proteins have a 33 amino acid insertion, lacking in mammals, and overexpression of chicken ANP32A (chANP32A) in human cells rescues efficient function of avian origin IAV polymerases^4^. Removal of the 33 amino acids from chANP32A prevents polymerase rescue, whilst conversely artificial insertion of the 33 amino acids into either huANP32A or B overcomes host restriction^4^. A naturally occurring splice variant of avian ANP32A lacks the first 4 amino acids of the 33 amino acid insertion, reducing the rescue efficiency of avian IAV polymerase in human cells^5^. This may be due to the disruption of a SUMOylation interaction motif, shown to enhance chANP32A’s interaction with IAV polymerase^6^. In human cells, both family members ANP32A and ANP32B (huANP32A/B) are utilised by human adapted IAV polymerases, and are thought to stimulate genome replication from the viral cRNA template, although the exact mechanism remains unclear^7^.

Here we demonstrate that avian ANP32B is evolutionarily distinct from mammalian and other ANP32Bs. We demonstrate that two amino acids differences, N129 and D130, in the LRR5 domain of chANP32B render it unable to interact with and support IAV polymerase function. We used CRISPR/Cas9 to remove the exon encoding the 33 amino acid insertion from chANP32A or to knockout the entire protein in chicken cells. Edited cells that expressed the short chANP32A isoform lacking the additional 33 amino acids supported mammalian-adapted but not avian IAV polymerase activity. Cells completely lacking chANP32A did not support either mammalian or avian IAV polymerase activity and were refractory to IAV infection. These results suggest a strategy to engineer IAV resistance in poultry through genetic deletion or single amino acid changes of the LRR domain of ANP32A protein.

## Results

### Phylogenetic analysis identifies that avian ANP32B is a paralog of mammalian ANP32B

To examine the relatedness of ANP32 proteins from different species, we constructed a phylogenetic tree using vertebrate ANP32 protein sequences using Drosophila mapmodulin protein as an outgroup. ANP32A and E homologues formed well-supported monophyletic clades which included multiple avian and mammalian species (Figure 1 & S1). Most vertebrate ANP32B proteins formed a monophyletic clade but this clade did not include avian ANP32B proteins. Rather, avian ANP32B proteins were strongly supported as members of a distinct ANP32C clade with ANP32C from *Xenopus* and unnamed predicted proteins (ANP32C) from non-placental mammals. This suggests that avian ANP32B and mammalian ANP32B are paralogues: birds have lost the protein orthologous to human ANP32B and eutherian mammals have lost the protein orthologous to avian ANP32B. Synteny provides further evidence to support the relationship between avian ANP32B and ANP32C genes as *Xenopus* ANP32C, avian ANP32B and marsupial ANP32C are all found adjacent to ZNF414 and MYO1F on their respective chromosomes (Figure S2).

**Figure 1.**
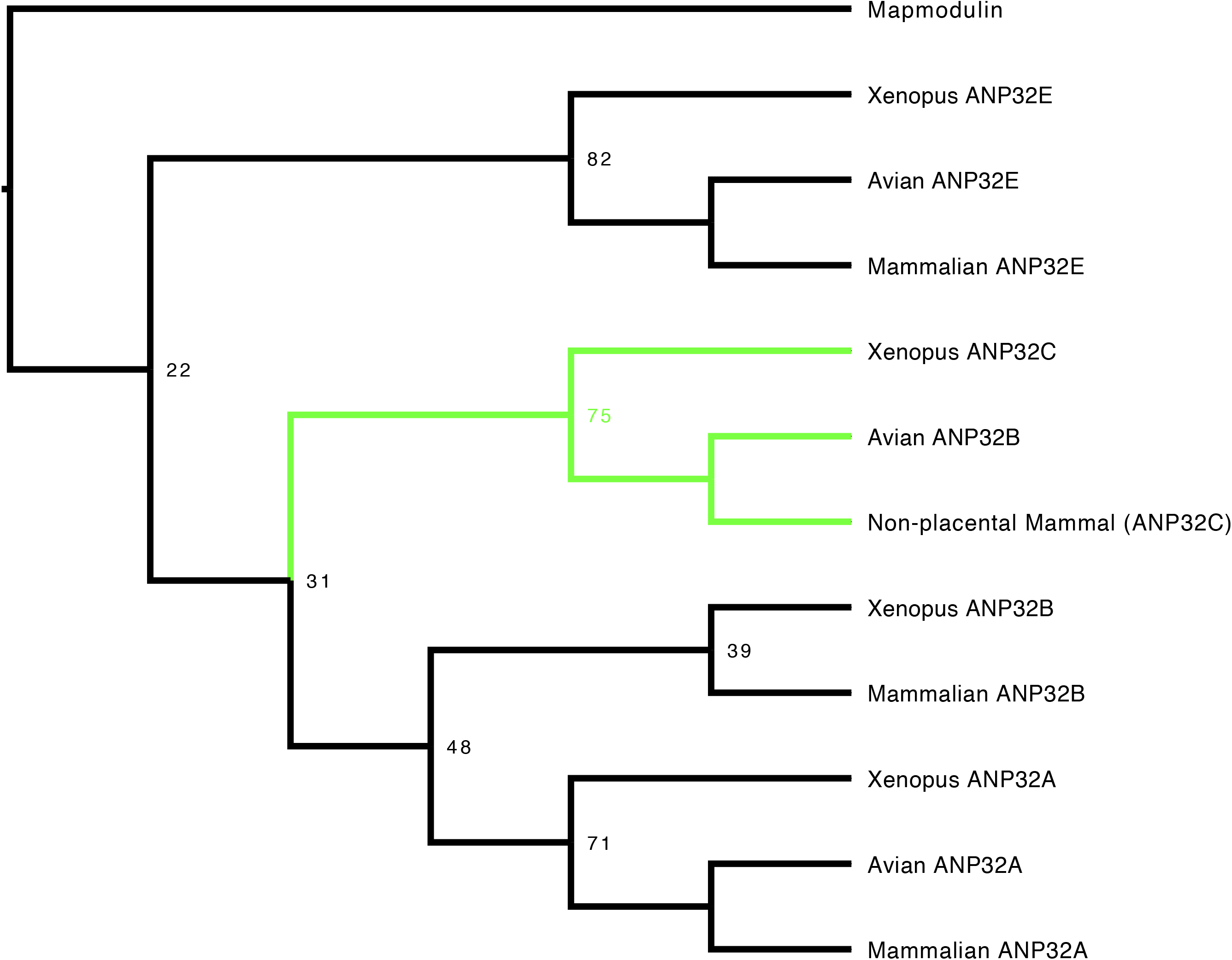
Phylogenetic and sequence analysis reveals avian ANP32B to be a paralog of mammalian ANP32B. The best maximum-likelihood tree was calculated from a set of ANP32 proteins with mapmodulin from *Drosophila melanogaster* as an outgroup using RAxML with 100 bootstraps. This figure is a cladogram showing the relationships between mammalian ANP32s, avian ANP32s and ANP32s from *Xenopus tropicalis*. Selected bootstrap values show the relationship between different ANP32 protein clades. Avian ANP32B clade is shown in green. The full tree is shown in Figure S1.

### Chicken ANP32B does not support lAV polymerase activity

We and others have previously shown that both human ANP32A and B proteins support activity of a human-adapted IAV polymerase in human cells ^4,7,8^. Using CRISPR/Cas9, we generated human eHAP1 cells that lacked expression of both human ANP32A and ANP32B protein (Staller et al. *in preparation*). In WT eHAP1 cells, human-adapted IAV polymerase (PB2 627K) was active, whereas avian polymerase (PB2 627E) was not. Exogenous expression of C-terminally FLAG-tagged chANP32A could rescue the activity of avian IAV polymerase whereas expression of chANP32B-FLAG, which naturally lacks the 33 amino acid insertion, did not (Figure 2a). In double knockout cells, neither human-adapted nor avian-origin polymerase were active. Expression of chANP32A-FLAG rescued activity of both polymerases but expression of chicken ANP32B-FLAG rescued neither, despite confirmation of robust expression by western blot (Figure 2b & c). This suggests that chicken ANP32B is not functional for IAV polymerase and that the IAV polymerase activity relies on ANP32A in chicken cells. To confirm this in chicken cells, we used CRISPR/Cas9 gene editing to generated chicken DF-1 cells which lacked chANP32B but retained chANP32A expression (DF-1 bKO Figure S3a). Wild type DF-1 cells had mRNAs for chANP32A, B and E (Figure S3d) and supported activity of avian IAV polymerase bearing either PB2 627E or 627K. Overexpression of chANP32B-FLAG did not affect activity (Figure 2d). DF-1 bKO cells also supported activity of both polymerases and again, exogenous expression of chANP32B had no effect. Since chicken cells lacking expression of chANP32B did not demonstrate any loss of IAV polymerase activity compared to WT, this implied that chANP32B is not functional for IAV polymerase and that IAV polymerase uses solely ANP32 family member A in chicken cells.

**Figure 2.**
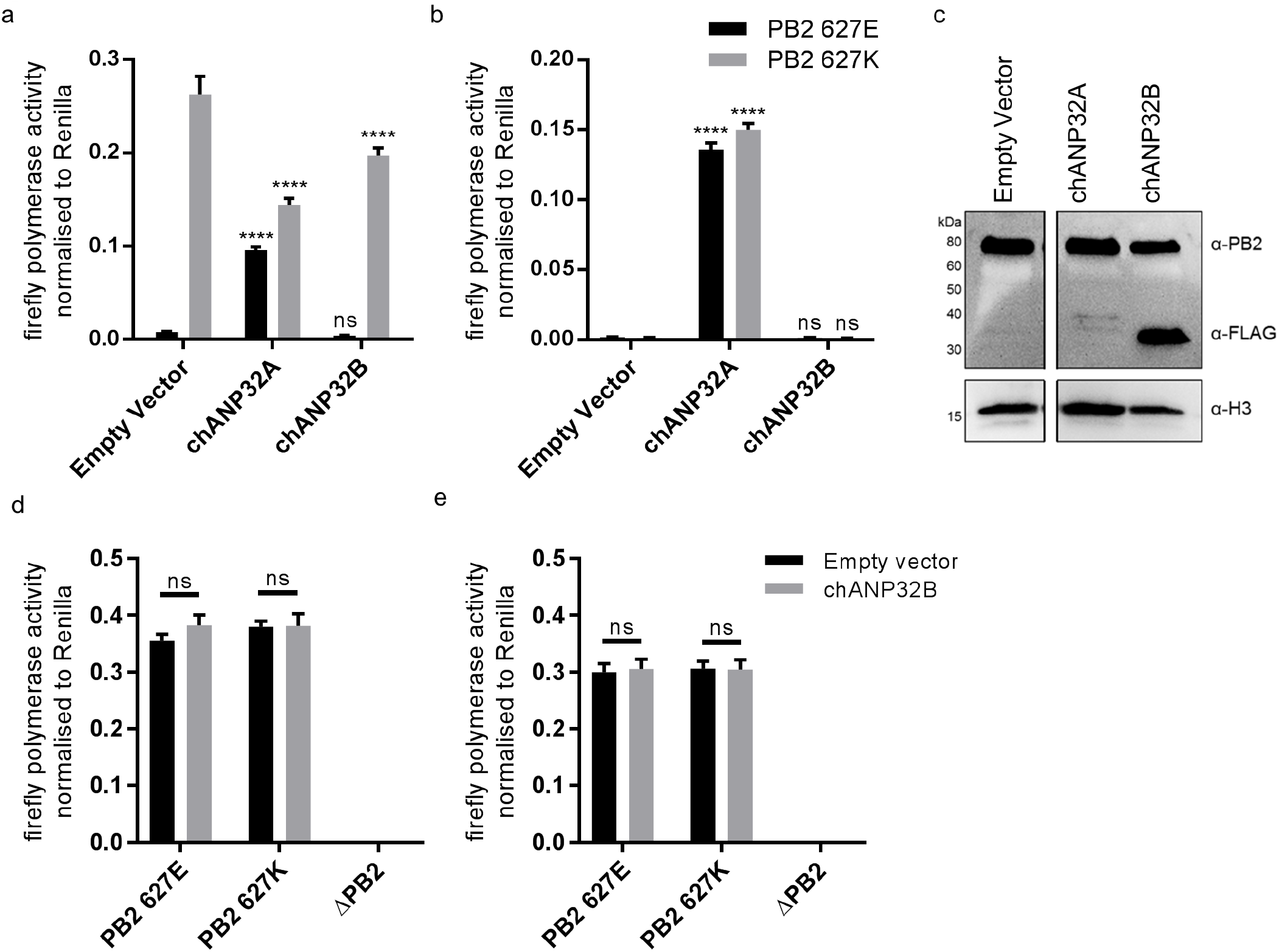
Chicken ANP32B is not functional for IAV polymerase. Cells were transfected with avian H5N1 50-92 polymerase (PB2 627E or 627K) together with NP, firefly minigenome reporter, *Renilla* expression control, either Empty vector (control) or ANP32 expression plasmid and incubated at 37°C for 24hours. **a.** Minigenome assay in human eHAP1 cells with coexpressed Empty vector, FLAG-tagged chANP32A or chANP32B. **b.** Minigenome assay in double knockout (dKO) eHAP1 cells. **c.** Western blot analysis of dKO eHAP1 cell minigenome assay confirming expression of PB2 and FLAG-tagged chANP32A and B. **d.** Minigenome assay in WT DF-1 cells with either co-expressed Empty vector or chANP32B. **e.** Minigenome assay in DF-1 ANP32B knockout (bKO) cells with either co-expressed Empty vector or chANP32B. Data shown are firefly activity normalised to *Renilla*, plotted as mean ±SEM. Two-way ANOVA with Dunnet’s multiple comparisons to Empty vector. ns= not significant, ****p<0.0001.

### Chicken cells lacking intact ANP32A do not support avian IAV polymerase activity

To investigate the function of ANP32A in chicken cells we utilised a cell type that is more amenable to genome editing and clonal growth. Primordial germ cells (PGCs) are the lineage restricted stem cells which form the gametes in the adult animal. PGCs from the chicken embryo can be easily isolated and cultured indefinitely in defined medium conditions^9,10^. Chicken PGCs can be edited using artificial sequence-specific nucleases and subsequently used to generated genome edited offspring^11,12^. Under appropriate *in vitro* conditions PGCs can acquire pluripotency and be subsequently differentiated into multiple cell types^13–16^. Chicken PGC cells were genome edited using CRISPR/Cas9 and appropriate guide RNAs to generate chANP32A knock-out cells (aKO) by targeted deletion of 8 nucleotides in exon 1. PGCs lacking the 33 amino acid insertion in chANP32A were generated using a pair of guide RNAs to remove exon 5 resulting in chicken cells with a mammalian-like ANP32A (Δ33) (Figure 3a). The precise deletions were confirmed by sequence analysis of genomic DNA (Figure S3c). We differentiated the edited chicken PGCs into fibroblast-like cells using serum induction with the aim of generating cell lines to test avian AIV polymerase activity (Figure S4). The predicted alterations of ANP32A protein in these cells were confirmed by western blot analysis of the PGC-derived fibroblast cells (Figure 3b). WT, Δ33, and aKO and PGC-derived cell lines were tested for the functional effects of alteration or loss of chANP32A expression on IAV polymerase activity measured by reconstituted minigenome assay. Both avian (PB2 627E) and human-adapted polymerase (PB2 627K) were active in WT fibroblast cells (Figure 3c). Removal of the 33 amino acids from ANP32A resulted in restriction of the 627E polymerase but not the 627K polymerase, mirroring the avian IAV polymerase phenotype observed in mammalian cells^4^. Both polymerases were restricted in cells lacking chANP32A (KO). Expression of exogenous chANP32A in Δ33 and aKO cells rescued avian IAV polymerase activity (Figure 3d & e) demonstrating the specificity of the genetic alterations. The lack of polymerase activity in the aKO PGC cell line supports the hypothesis that, in the absence of chANP32A, the remaining ANP family members including chANP32B or chANP32E could not support IAV polymerase activity in chicken cells, even though ANP32B and E mRNAs were readily detected in both DF-1 and PGC cells (Figure S3d & e).

**Figure 3.**
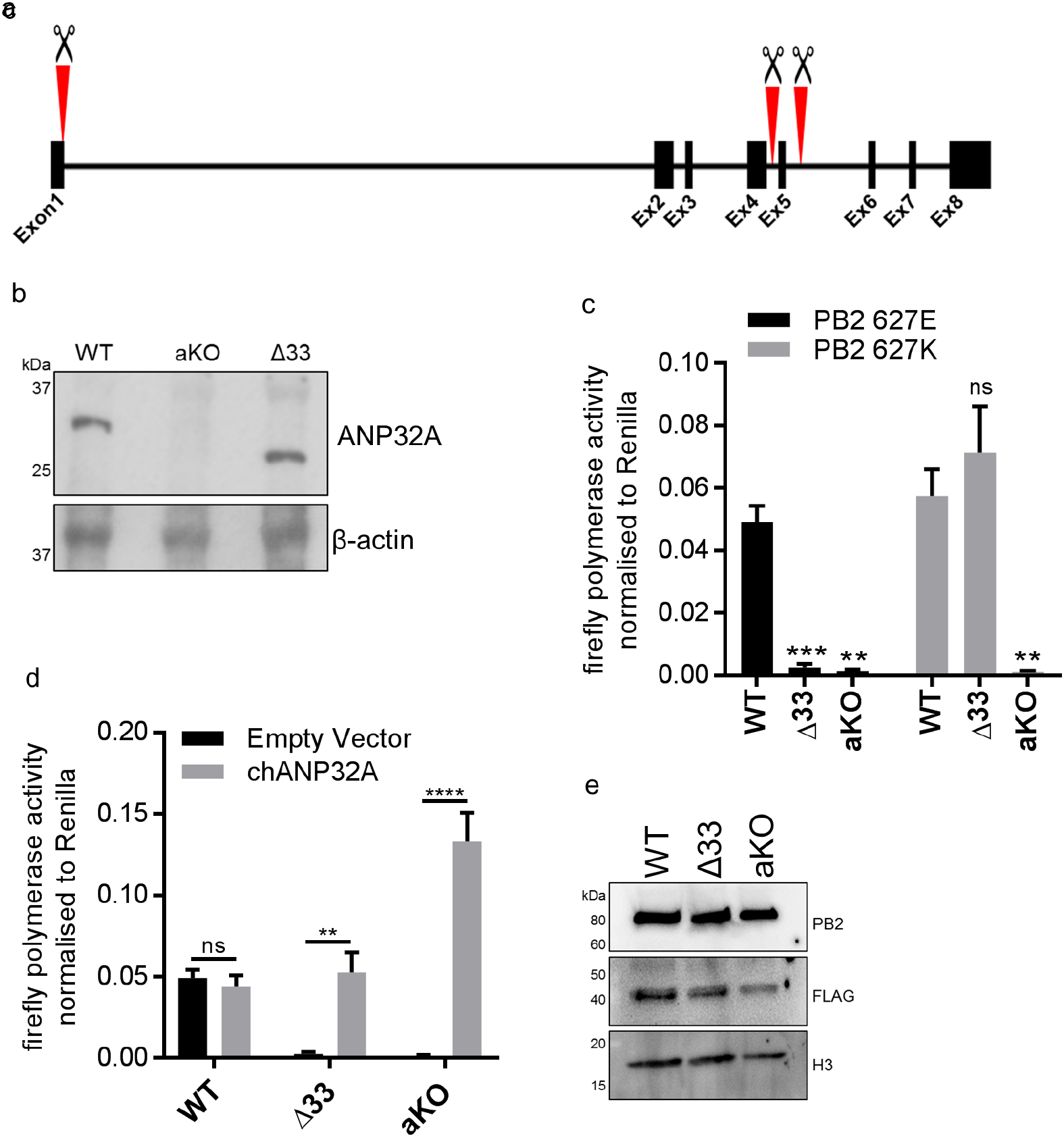
Chicken PGC derived fibroblast cells lacking ANP32A or the 33 amino acid insertion do not support avian IAV polymerase activity. **a.** Schematic of CRISPR/Cas9 RNA guide targets used to generate aKO (exon1) and Δ33 (exon 5) PGC cell lines. **b.** Western blot analysis of ANP32A and β-actin expression in WT, KO and Δ33 PGC-derived fibroblast cells. **c.** Minigenome assay in WT, Δ33 or aKO PGC derived fibroblast cells with either PB2627E (black) or 627K (grey) polymerase derived from avian H5N1 50-92 virus. **d.** Minigenome assay in WT, Δ33 or aKO cells with avian H5N1 50-92 PB2 627E polymerase co-transfected with Empty vector (black) or FLAG-tagged chANP32A (grey). **e.** Western blot analysis of PB2, FLAG and Histone 3. Data shown are firefly activity normalised to *Renilla*, plotted as mean ±SEM. Two-way ANOVA with Dunnet’s multiple comparisons to WT. ns= not significant, **P < 0.01, ***p<0.001, ****p<0.0001.

### Functional differences between chicken ANP32A and ANP32B map to the LRR domain sequence

ANP32 proteins share a common domain organization in which an N terminal domain consisting of 5 consecutive leucine rich repeats (LRR 1-5) is followed by a cap and central domain and a C terminal low complexity acidic region (LCAR). In avian ANP32A proteins (except some flightless birds) a sequence duplication, derived in part from nucleotides that encode 27 amino acids (149-175), has resulted in an additional exon and an insertion of up to 33 amino acids between the central domain and the LCAR (Figure 4a). We previously showed that insertion of the 33 amino acids from the central domain of chANP32A into the equivalent region of the human ANP32A or huANP32B proteins conferred the ability to rescue the activity of a restricted avian IAV polymerase in human cells. The equivalent 33 amino acid insertion into chANP32B (chANP32B^33^) did not support avian IAV polymerase activity (Figure 4b). In order to ascertain the domains of chANP32B that rendered it non-functional for IAV polymerase activity, we generated chimeric constructs between human and chicken ANP32B. To measure the rescue of avian IAV polymerase in human 293T cells, all chimeric constructs had the 33 amino acid sequence derived from chANP32A inserted between the LRR and LCAR domains. Western blot analysis and immunofluorescence confirmed that all chimeric constructs were expressed and localized to the cell nucleus as for the wild type ANP32 proteins. (Figure 4b & S5). Swapping the LCAR domain of chANP32B into huANP32B^33^ did not prevent the rescue of avian IAV polymerase (huANP32B^33^_LCAR_). Introduction of the central domain of chANP32B into huANP32B (huANP32B^33^_CENT_) significantly reduced rescue efficiency and swapping the LRR domain of chANP32B (huANP32B^33^_LRR_) rendered the protein non-functional to avian IAV polymerase (Figure 4b). By sequential swapping of each LRR repeat, the 5^th^ LRR of chANP32B was found to be the domain that prevented rescue of avian IAV polymerase (Figure 4b). The fifth LRR contains five amino acid differences between human and chicken ANP32B, highlighted on the crystal structure of huANP32A, plus an additional one difference to chANP32A (Figure 4d & S6). Swapping chANP32B’s fifth LRR into chANP32A also prevented rescue of avian IAV polymerase activity in human cells (chANP32A_LRR5_) (Figure 4c). Introduction of the single amino acid changes derived from the chANP32B LRR5 sequence into chANP32A revealed that mutations N129I and D130N significantly reduced the ability of chANP32A to rescue avian IAV polymerase activity in human cells (Figure 4c). Minigenome assays with co-expressed chANP32A or chANP32A_N129I_ in aKO chicken fibroblast cells confirmed that the 129I mutation significantly reduced the ability of chANP32A to support avian-origin (PB2 627E) or human-adapted (PB2 627K) IAV polymerase activity (Figure 4e).

**Figure 4.**
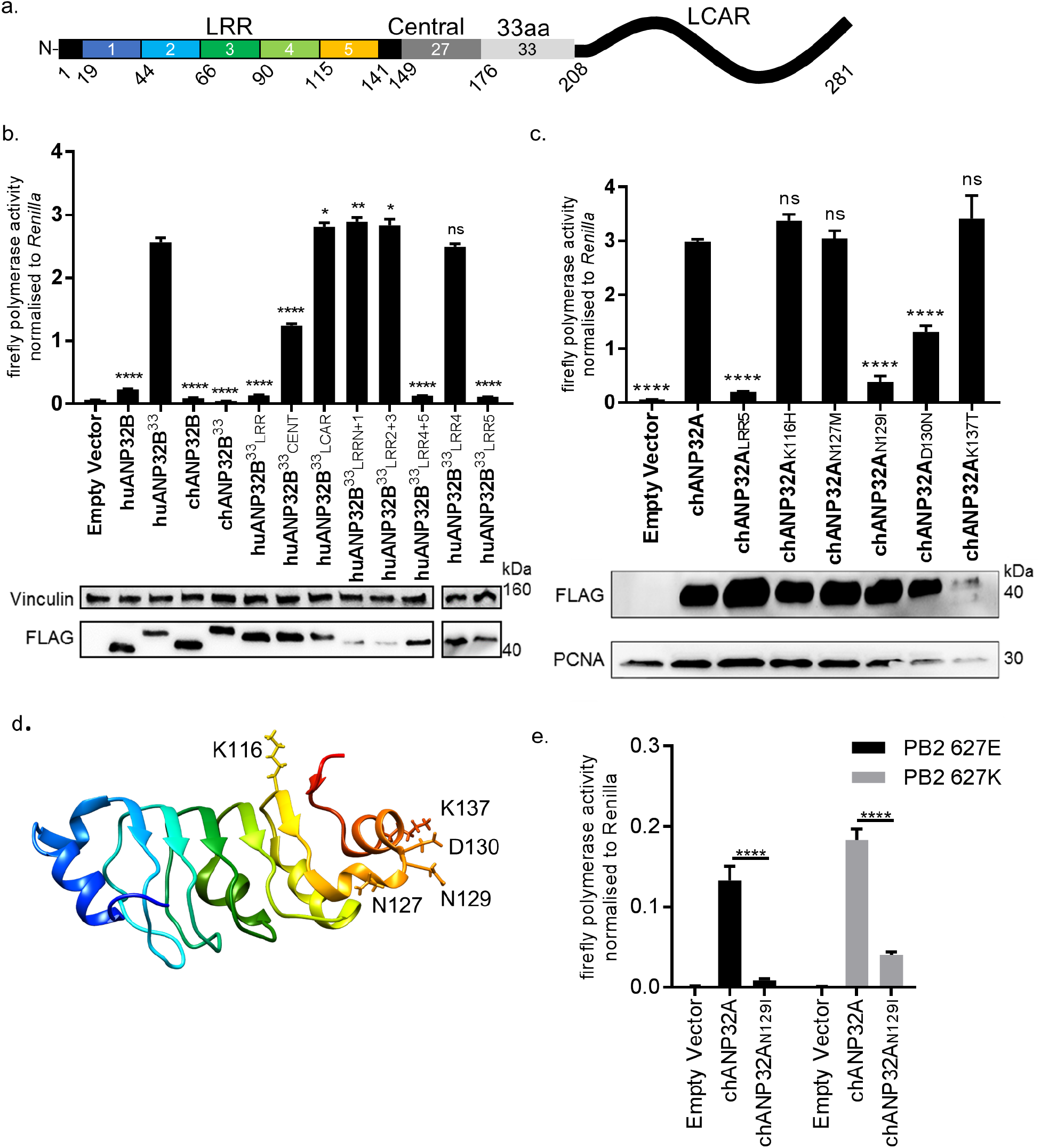
Lack of functional support for IAV polymerase by chicken ANP32B maps to differences in LRR5 domain. **a.** Schematic of chicken ANP32A protein highlighting the different domains and LRR sequences (LRR 1-5). **b.** Human 293T cells were transfected with avian H5N1 50-92 polymerase (PB2 627E) together with NP, pHOM1-firefly minigenome reporter, *Renilla* expression control, either Empty vector or FLAG-tagged ANP32 expression plasmid and incubated at 37°C for 24hours. Western blot analysis shown below (FLAG and Vinculin). **c.** 293T minigenome assay in 293T cells (PB2 627E) with FLAG-tagged WT or mutant chANP32A expression plasmids with associated western blot (FLAG and PCNA). **d.** huANP32A crystal structure (PDB 4X05) with residues K116, N127, N129, D130 and K137 highlighted using UCSF Chimera ^25^. **e.** Minigenome assay of avian H5N1 50-92 polymerase with either PB2 627E or 627K in PGC-derived fibroblast aKO cells, together with co-expressed Empty vector, chANP32A or chANP32A_N129I_. Data shown are firefly activity normalised to *Renilla*, plotted as mean ±SEM. One-way ANOVA with Tukey’s comparison to chANP32A (b&c) or two-way ANOVA with Dunnet’s multiple comparisons to chANP32A (e). ns= not significant, *p<0.05, **P < 0.01, ****P < 0.0001.

### Sequence of amino acids 149-175 of the central domain of chANP32A are required to support activity of both avian and human-adapted IAV polymerase

As chANP32A KO PGC-derived fibroblast cells did not support of IAV polymerase despite expressing chANP32B, we were able to use these cells to understand in more detail the sequences in chANP32A required for IAV polymerase activity. The results above showed that the 33 amino acid insertion, fifth LRR and central domain are important for the ability of chANP32A to support function of avian IAV polymerase. We performed the minigenome assay in aKO cells with polymerases containing either PB2 627E and 627K with co-expression of further chANP32 mutants including: chANP32A in which the 27 amino acids in the central domain preceding the 33 amino acid insertion were scrambled (chANP32A_scri49-17s_) or chANP32A with the 33 amino acid insertion scrambled (chANP32A_scr176-208_) (Figure 5a). Both mutants were expressed and localized to the nucleus (Figure 5c and S5). The first mutant, chANP32A_scr149-175_, did not support either PB2 627E or 627K polymerase, suggesting the sequence of the central domain is important for function of IAV polymerase. The second mutant, chANP32A_scr176-208_, only supported PB2 627K function, confirming that the sequence of the 33 amino acid insertion, not just the extended length is required for avian IAV polymerase (PB2 627E) (Figure 5b).

**Figure 5.**
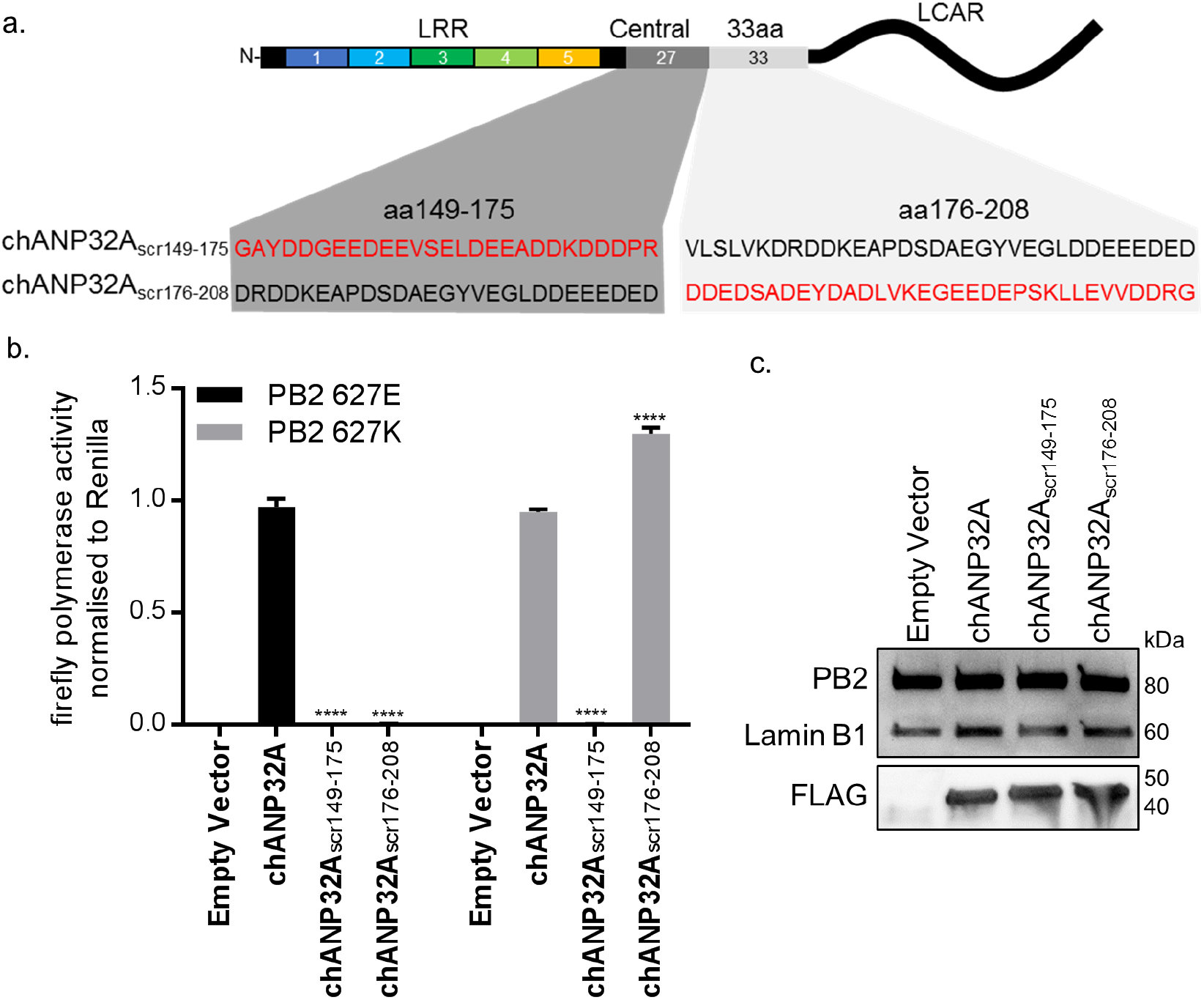
Sequence of amino acids 149-175 of the central domain of chANP32A are required to support activity of both avian and human-adapted IAV polymerase. **a.** Schematic of chANP32A showing the sequence of amino acids in the central domain (149-175 or 33 amino acid insertion (176-208) and the randomly scrambled sequence in red. **b.** Minigenome assay of avian H5N1 50-92 polymerase with either PB2 627E or 627K in PGC-derived fibroblast aKO cells with co-expressed Empty plasmid or FLAG-tagged WT chANP32A, chANP32A_scr149-175_ or chANP32A_scr176-208_ expression plasmids. **c.** Western blot analysis of PB2 (627E), lamin B1 and FLAG. Data shown are firefly activity normalised to *Renilla*, plotted as mean ±SEM. Two-way ANOVA with Dunnet’s multiple comparisons to chANP32A. ns= not significant, ****P < 0.0001.

### A single amino acid difference between chANP32B and chANP32A abrogates binding of chANP32A to IAV polymerase

An interaction between ANP32A and IAV polymerase was demonstrated previously that is dependent on the presence of all three polymerase subunits (Mistry et al. *in preparation* & ^5,6^). To examine the interaction between IAV polymerase and chANP32 proteins we employed a split luciferase complementation assay as a quantitative measure of binding ^17,18^. The C-terminus of the PB1 subunit of avian origin IAV polymerase was fused with one half of *gaussia* luciferase (PB1^luc1^) and the C-terminus of chicken ANP32A or B with the second half (chANP32A^luc2^ and chANP32B^luc2^) (Figure 6a). Reconstitution of PB1^luc1^, PB2 and PA together with chANP32A^luc2^ in human 293T cells gave a strong Normalised Luciferase Ratio (NLR) (Figure S6a) with polymerases containing either PB2 627E or 627K (Figure 6b). Luciferase complementation was significantly less between polymerase and chANP32B^luc2^, and even insertion of the 33 amino acids from chANP32A did not restore the signal (chANP32B_33_^luc2^) (Figure 6b). When chANP32A carried the single N129I mutation (chANP32A_N1291_^luc2^), luciferase complementation was reduced 22-fold for PB2 627E polymerase and 52-fold forPB2 627K polymerase (Figure 6c & S7). These results suggest that the loss of support of polymerase function by chANP32A_N129I_ was due to a disruption of binding to IAV polymerase.

ANP32A proteins bind to histones as part of their role in chromatin regulation ^19^. To measure if the mutation N129I had any effect on this cellular interaction, we generated expression plasmids that encoded human histone 4 with luc1 fused to the C-terminus (H4^luc1^) and histone 3.1 with luc2 fused to the C terminus (H3^luc2^). As expected, H4^luc1^ and H3^luc2^ generated a strong NLR, reflecting their interaction in the nucleosome ^20^. The ability of chANP32A to bind histone 4 was not impaired by mutation N129I, suggesting chANP32N129I was not altered in this cellular role, despite abrogation of its support of IAV polymerase (Figure 6d).

**Figure 6.**
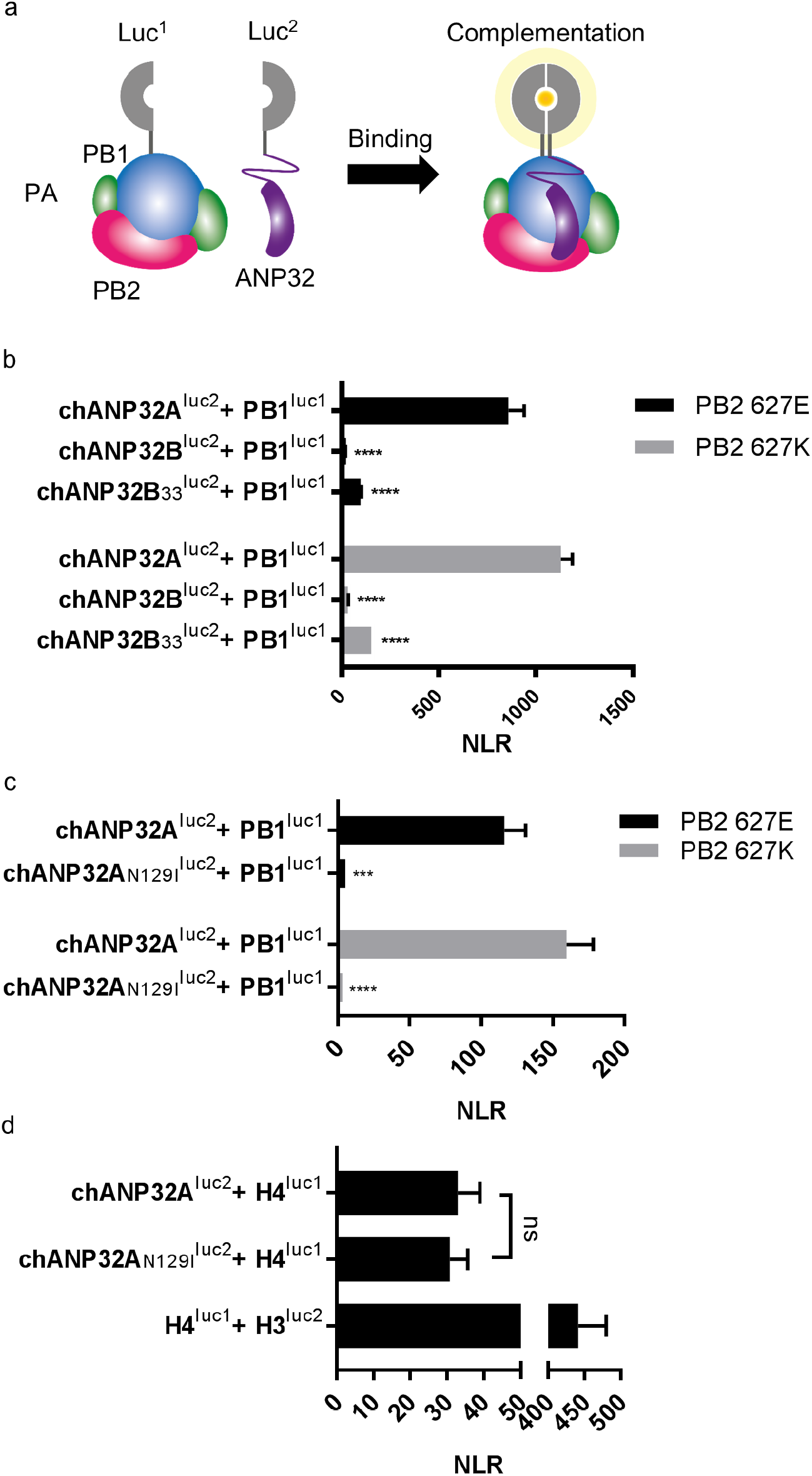
A single amino acid change (N129I) derived from chANP32B disrupts chANP32A support of influenza polymerase activity by abrogating binding to IAV polymerase. **a.** Diagram of the split *Gaussia* luciferase system, demonstrating how ANP32 fused to luciferase fragment luc2 may bind to polymerase containing PB1 fused to luciferase fragment luc1 and complement full luciferase, which then reacts with substrate to generate a measurable bioluminescent signal. **b.** Human 293T cells were transfected with PB1 fused to luc1 (PB1^luc1^), PB2 (627E or K), PA and either chANP32A, chANP32B or chANP32B33 fused to luc2 (control wells were transfected with all components but with unfused PB1 and luc1 or chANP32 and luc2). **c.** As (b) but with either chANP32A^luc2^ or chANP32A_N129I_^luc2^. **d.** 293T cells transfected with either chANP32A^luc2^ or chANP32A_N129I_^luc2^ and histone 4 fused to luc1 (or with unfused controls) or with H4luc1 and histone 3 fused to luc2. All data are Normalised Luciferase Ratio (Figure S5). One-way ANOVA (d) or two-way ANOVA with Dunnet’s multiple comparisons to chANP32A (b&c). ns= not significant, ****P < 0.0001.

### Viral replication is abrogated in chicken cells lacking ANP32A

The data above suggest that chANP32B cannot substitute for chANP32A in support of IAV polymerase in chicken cells. Since chicken cells that completely lack expression of chANP32A show no polymerase activity in the minigenome assay, they might be refractory to IAV infection. Multi-cycle growth kinetics of recombinant influenza A viruses were measured in WT, Δ33 and aKO PGC-derived fibroblast cells (Figure 7). To ensure robust infection, recombinant viruses were generated carrying H1N1 vaccine strain PR8 haemagglutinin (HA), neuraminidase (NA) and M genes; this also mitigated the risks of working with avian influenza viruses with novel antigenicity. Infectious titres of recombinant virus with internal genes of avian H5N1 virus A/England/1991/50-92 were reduced almost 3000-fold at 24 hours post infection in absence of chANP32A protein (Figure 7a). Similarly, a recombinant virus with internal genes from avian H9N2 virus A/duck/Belgium/24311/12 also replicated poorly in aKO cells (Figure 7b). An isogenic pair of viruses with internal genes from the H7N9 virus A/Anhui/1/2013, carrying either PB2 627E or 627K, replicated efficiently in WT fibroblasts, but not in aKO cells where peak viral titres were more than 200-fold less (Figure 7c & d). Comparison of viral titres in supernatant recovered at 12 hours post infection and later time points by two-way ANOVA revealed that there was no statistically significant increase over time in viral titres released from aKO cells, in contrast with WT cells where viral titres were significantly amplified at later time points (Figure 7). In conclusion, PGC derived fibroblast cells lacking chANP32A were effectively resistant to IAV replication.

**Figure 7.**
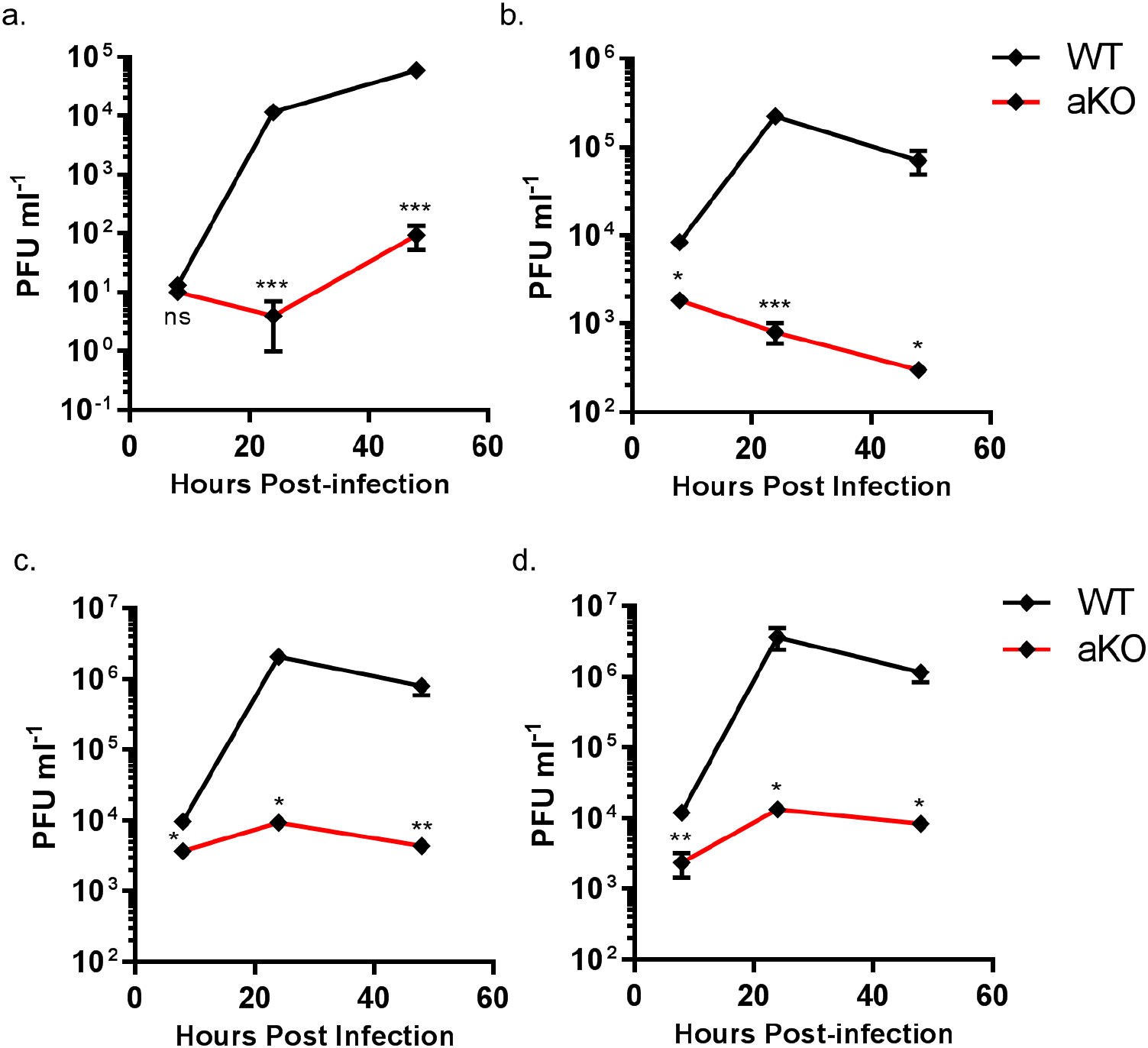
Viral replication is abrogated in chicken PGC fibroblast cells lacking ANP32A. WT (black lines) or aKO (red lines) PGC-derived fibroblast cells were infected with reassortant viruses (containing PR8 HA, NA and M genes and internal genes from avian IAVs), incubated at 37°C in the presence of trypsin, and cell supernatants harvested at described time-points and PFU ml^−1^ measured by plaque assay on MDCK cells. **a.** H5N1 50-92 (MOI 0.001). **b.** H9N2 Belgium (MOI 0.005). H7N9 Anhui PB2 627E **(c)** or PB2 627K **(d)** (MOI 0.005). Multiple t-tests with Holm-Sidak comparison. ns= not significant, *p<0.05, **P<0.01, ***p<0.001.

## Discussion

We show that avian origin IAV polymerases rely exclusively on chicken ANP32A family member for their replication, because they are unable to co-opt chicken ANP32B. We found avian ANP32B proteins formed a separate phylogenetic group from other ANP32Bs and were more closely related to ANP32C proteins of other species than to mammalian ANP32B members (Figure 1 & S1). Synteny demonstrated that ANP32C was present in coelacanth, amphibians and non-placental mammals and that this locus was identical to ANP32B in birds (Figure S2). ANP32C has been lost in placental mammals. Human ANP32C is an intronless gene and has been suggested to be a pseudogene derived from ANP32A^19^. There was no evidence of a mammalian ANP32B equivalent in birds. We propose that avian ANP32Bs should be renamed as ANP32Cs.

Chicken ANP32B could not support influenza polymerase function due to an amino acid difference in LRR5 at residue 129 that adversely affected the interaction between chANP32B and influenza polymerase (Figure 6). Other avian ANP32B proteins, including those of duck and turkey, carry isoleucine at residue 129 suggesting that our findings may also be applicable to other avian hosts (Figure S6). The replacement of the exposed polar residue, asparagine (N129) with the hydrophobic isoleucine (I) may have led to the disruption of a key electrostatic interaction between ANP32A and the virus polymerase complex. In addition to the residue 129I, the central domain (amino acids 141-175) of chANP32B also contributed to its poor efficiency at rescuing avian IAV polymerase function in human cells (Figure 4). This, together with the observation that scrambling amino acids 149-175 in chANP32A prevented both human-adapted and avian IAV polymerase function (Figure 4) suggests that LRR5 and the central domain of ANP32A are crucial to IAV polymerase function. The observation that scrambling the 33-amino acid insertion prevented avian IAV polymerase rescue (Figure 5) is consistent with results from by Domingues and Hale and Baker and colleagues which showed that the SUMO Interaction motif (SIM)-like sequence present in the 33 amino acid insertion (VLSLV), was required for strong binding to both 627E and 627K polymerase and its deletion or mutation decreased its ability to support avian IAV polymerase activity in human cells^5,6^. Understanding the domains important to binding and function may help us understand the mechanism by which ANP32A or B support IAV polymerase which is still not fully elucidated^7^.

Chicken cells lacking ANP32A did not support activity of avian and human-adapted IAV polymerase in minigenome assay (Figure 2). Very low viral titres were recovered following infection in the aKO fibroblast cells (Figure 7). IAV polymerase may function, albeit inefficiently, in the absence of chANP32A in the context of virus infection. Other viral products present during virus infection such as NEP may partly compensate for the block in replication in cells that lack chANP32A. Indeed NEP expression has been reported to rescue avian polymerase replication in human cells^21^. Nonetheless the very low level of replication observed in the chicken cells that lack chANP32A *in vitro* implies that *in vivo*, chickens that do not express ANP32A may be resistant to infection.

The use of the PGC-derived chicken cells to investigate a host factor essential for virus raises the possibility of generating genome-edited chicken models resistant or resilient to infection. Chicken PGCs can be efficiently genome-edited to generate specific haplotypes^22^. Our novel method of chicken PGC differentiation into fibroblast-like cells enabled robust testing of a defined genotype, and will permit future investigation of other host genetic factors identified through forward genetic screens and suspected to play important roles in virus infections^23,24^.

In summary, we provide evidence that specific domains of ANP32 proteins are important for the function of IAV polymerases and describe a lack of redundancy in the involvement of ANP32 family members to support IAV polymerase complex in chicken cells that is determined by the variation in ANP32 protein sequences. These data may aid in the design of novel small molecule inhibitors that disrupt the ANP32-polymerase interface and form the basis of a potential pathway for the generation of influenza virus-resistant animals.

## Acknowledgments

JSL, CR, SG, MAS, HMS and WSB were supported by Biotechnology and Biological Sciences Research Council (BBSRC) via Strategic LoLa grant BB/K002465/1 “Developing Rapid Responses to Emerging Virus Infections of Poultry (DRREVIP)”. AI-A was funded by a Principal’s Career Development PhD Scholarship from the University of Edinburgh. BM was supported by the Wellcome Trust. DHG, JS and WSB were supported by grant 205100 from the Wellcome Trust. ES was supported by an Imperial College President’s Scholarship. HS was funded from BBSRC grants BB/R007292/1 and BBS/E/I/00007034. HMS and MJM were supported by Institute Strategic Grant Funding from the BBSRC (BB/P013732/1 and BB/P013759/1).

## Author contributions

JSL, AI-A, SG, HS, MAS, HMS, MJM and WSB designed the research. JSL, AI-A, BM, JS, CR performed experiments. ES and HS contributed new reagents. JSL, AI-A, DHG, MJM analysed data. JSL, AI-A, DHG, SG, HS, MAS, HMS, MJM and WSB wrote the paper.

## Methods

### Animal use

The GFP+ PGCs used in the experiments were obtained by crossing the Roslin Green (ubiquitous GFP) line of transgenic chickens with a flock of commercial Hyline layer hens maintained at the Roslin Institute to produce heterozygous fertile eggs for PGC derivations^27^. Commercial and transgenic chicken lines were maintained and bred under UK Home Office License. All experiments were performed in accordance with relevant UK Home Office guidelines and regulations. The experimental protocol and studies were reviewed by the Roslin Institute Animal Welfare and Ethical Review Board (AWERB) Committee.

### Plasmid constructs

ANP32A guide RNAs (gRNA) were designed using CHOPCHOP gRNA web tool (http://chopchop.cbu.uib.no/)^28,29^. gRNA 5′-CGGCCATGGACATGAAGAAA-3′ targeting ANP32A exon1, and gRNAs:5′-AGCTGGAAGCAATATGTACT-3′ and 5′-CATTCCCCTCGCTCCTTCAA-3′ targeting either side of exon 5 (Δ33 PGC cells) were cloned into pSpCas9(BB)-2A-Puro (pX459 v2.0; a gift from Dr. Feng Zhang) using materials and methods described by previously^30^. For DF-1 ANP32B gRNAs, the guides 5′- TTCAGATGATGGGAAGATCG-3′ and 5′-GGTTCTCAAAATCTGAAGAG-3′ were cloned into the double “nickase” vectors pSpCas9n(BB)-2A-GFP (pX461) and pSpCas9n(BB)-2A-Puro (pX462) respectively^30,31^.

*Gaussia* luc1 and luc2 were generated by gene synthesis (GeneArt, ThermoFisher) using the sequence previously described^18^. *Homo sapiens* Histone 4 (NP_003533.1) and 3.1 (NP_003520.1) were generated gene synthesis (GeneArt, ThermoFisher). Luc1 or luc2 were added to the C-termini of ANP32, PB1, H4 or H3.1 using the linker sequence, AAAGGGGSGGGGS, by overlapping PCR. The 33 amino acid insertion was added to huANP32B after residue 173 and to chANP32B after residue 181 (preserving an acid region before SIM motif^6^). The LRR (amino acids 1-149), central domain (amino acids 150-175) or LCAR (amino acids 176-262) from chANP32B were swapped into huANP32B33 to generate chimeric constructs. ANP32 constructs were made by overlapping PCR or by gene synthesis (GeneArt, ThermoFisher) with either a FLAG tagged fused to the C-terminus with a GSG linker or to mCherry with a GSGGGSGG linker.

### Cells and cell culture

Human embryonic kidney (293T) (ATCC), human lung adenocarcinoma epithelial cells (A549) (ATCC) and Madin-Darby canine kidney (MDCK) cells (ATCC) were maintained in cell culture media (Dulbecco’s modified Eagle’s medium (DMEM; Invitrogen) supplemented with 10% fetal calf serum (FCS) (Biosera), 1% non-essential amino acids (NEAA) and with 1% penicillin-streptomycin (Invitrogen)) and maintained at 37 °C in a 5% CO2 atmosphere. Human eHAP1 cells were cultured in Iscove’s Modified Dulbecco’s Medium (IMDM) supplemented with 10% fetal bovine serum (FBS), 1% NEAAs, and 1% penicillin/streptomycin. Chicken fibroblast (DF-1) (ATCC) cells were maintained in DF-1 cell culture media (DMEM supplemented with 10% FCS, 5% tryptose phosphate broth (Sigma-Aldrich), 1% NEAAs and 1% penicillin-streptomycin and maintained at 39 °C in a 5% CO2 atmosphere.

### PGC, DF-1 and eHAP1 cell line generation

PGCs were derived and cultured in FAOT medium as previously described^10^. PGCs were transiently transfected with 1.5 μg of PX459 V2.0 vector using Lipofectamine 2000 (Invitrogen) and treated with puromycin as previously described^22^. Subsequently, single cell cultures of puromycin-resistant cells were established to generate clonal populations for downstream experiments as previously described in ^22^. To identify an ANP32A Δ33 PGC cell line, PCR products were directly sequenced using PCR primers to analyse mutation genotypes of isolated single cell clones. To identify an ANP32A KO PGC cell line, PCR products were cloned into pGEM-T Easy vector (Promega) and sequenced using T7 promoter forward primer by Sanger sequencing. DF-1 cells were transfected with the described CRISPR/Cas9 constructs using Lipofectamine 2000 (Invitrogen) and subject to puromycin selection. Single cell clones were expanded and analysed by PCR of genomic DNA and Sanger sequencing using primers (5′- TTTTTGCTTACATCTGAGGGC-3′, 5′-CCTCCGCAGTTATCAGGTTAGT-3′) for ANP32A exon1, (5′- GCTCCCTGGTCTGCTAGTTAT-3′, 5′-GGTCTACGCAACCACACATAC-3′) for ANP32A exon 5 and (5′- CCCTTAAGGTGAGCACAGGG-3′, 5′-AACATAGCACCACTCCCAGC-3′) for ANP32B exon2. eHAP1 dKO cells were generated as described (Staller et al. *in preparation)*.

### Differentiation of PGCs into adherent fibroblast-like cells (PGC derived fibroblasts)

PGCs were cultured in 500 μl of high calcium FAOT medium containing 1.8 mM CaCl2 in fibronectin-coated wells (24-well plate) for 48 hours (Figure S3)^10^. Subsequently, PGCs were transferred into PGC fibroblast medium and then refreshed every 48 hours by removing and replacing with 300 μl of PGC fibroblast cell culture medium. Adherent fibroblast-like cells were observed within 72 hours. Cells were then refed every two days and split 1:4 every four days. PGC fibroblast cell cultures were expanded to 85-90% confluency in 24-well plates before using for transfection, infection or western blot analysis. PGC fibroblast cells were maintained in cell culture media (Knockout DMEM (10829018, Gibco) with 10% ES grade FBS (16141061, Invitrogen), 1 % chicken serum (Biosera), 0.1% 100xNEAA (Gibco), 0.1% Pyruvate (11360070, Gibco), 0.1% 100xGlutamax (Gibco: 35050-038), 0.5mg ml^−1^ ovotransferin (C7786, Sigma)) and 1% penicillin-streptomycin at 37°C with 5% CO2.

### Influenza A virus infection

Recombinant influenza A PR8 (A/PR/8/34 (H1N1)) 3:5 reassortant viruses (PR8 HA, NA and M genes with PB1, PB2, PA, NP and NS genes from either A/duck/Belgium/24311/12 (H9N2) or A/Anhui/1/13 (H7N9) were generated by reverse genetics at The Pirbright Institute, UK. Reverse genetics virus rescue was performed by transfection of Human Embryonic Kidney (HEK) 293T cells (ATCC) with eight bi-directional PHW2000 plasmids containing the appropriate influenza A virus segments and coculture in Madin-Darby Canine Kidney (MDCK) cells (ATCC) with addition of 2μg ml^−1^ of TPCK treated Trypsin (Sigma-Aldrich). Rescued viruses were passaged once in embryonated hen’s eggs to generate working stocks. Recombinant PR8 3:5 reassortant 50-92 (A/turkey/England/50-92/1991 (H5N1) was described previously^4^.

Virus was diluted in Knockout DMEM and incubated on PGC fibroblast cells for 1 h at 37 °C (MOI as indicated in the relevant figure legends) and replaced with infection media (Knockout DMEM (10829018, Gibco), 0.14% BSA and 1 μg ml^−1^ TPCK trypsin (Sigma-Aldrich)). Cell supernatants were harvested and stored at −80°C. Infectious titres were determined by plaque assay on MDCK cells.

### Minigenome assay

Influenza polymerase activity was measured by use of a minigenome reporter which contains the firefly luciferase gene flanked by the non-coding regions of the influenza NS gene segment, transcribed from a species-specific polI plasmid with a mouse terminator sequence. The human and chicken polI minigenomes (pHOM1-Firefly and pCOM1-Firefly) are described previously^32^. pCAGGS expression plasmids encoding each polymerase component and NP for 50-92 (H5N1 A/Turkey/England/50–92/91) are described previously^33^. To measure influenza polymerase activity, 293T cells were transfected in 48-well plates with pCAGGS plasmids encoding the PB1 (20 ng), PB2 (20 ng), PA (10 ng) and NP (40 ng) proteins, together with 20 ng species-specific minigenome reporter, either Empty pCAGGS or pCAGGS expressing ANP32 (50ng) and, as an internal control, 10 ng Renilla luciferase expression plasmid (pCAGGS-Renilla), using Lipofectamine 3000 transfection reagent (Invitrogen) according to manufacturers’ instructions. DF-1 and PGC fibroblast cells were transfected as 293T cells but with twice the concentration of DNA. Cells were incubated at 37°C. 20-24 h after transfection, cells were lysed with 50 μl of passive lysis buffer (Promega), and firefly and Renilla luciferase bioluminescence was measured using a Dual-luciferase system (Promega) with a FLUOstar Omega plate reader (BMG Labtech).

### Split luciferase assay

293T cells were transfected with PB1^luc1^ (25ng), either PB2 627E or PB2 627K (25ng), PA (12.5ng) and chANP32A^luc2^, chANP32AN129I^luc2^, chANP32B^luc2^ or chANP32B33`^uc2^ (12.5ng). For split luciferase assays measuring histone interaction, 50ng of either chANP32A^luc2^, chANP32AN129I^luc2^, H4^luc1^ or H3^luc2^ were transfected into 293T cells. Control samples assessed the interaction between H4 or PB1^luc1^ and an untagged luc2 construct or the appropriate ANP32A^luc2^ construct and an untagged luc1 construct. All other components transfected into control samples remained consistent with those transfected in with the interacting proteins of interest. 24 hours after transfection, cells were lysed in 50ul Renilla lysis buffer (Promega) for one hour at room temperature. *Gaussia* luciferase activity was then measured from 10ul of lysate using the Renilla luciferase kit (Promega) with a FLUOstar Omega plate reader (BMG Labtech). Normalised luminescence ratios were calculated by dividing the luminescence measured from the interacting partners by the sum of the interaction measured from the two controls for each sample (Figure S5a) as previously described^18^.

### Immunoblot analysis

For analysis of PGC derived fibroblasts (Figure 3b), at least 300,000 cells were lysed in 60 μl of 1X RIPA lysis buffer (sc-24948, Santa Cruz Biotechnology) according to the manufacturer’s instruction. Protein concentration was determined using the Bradford method with the Quick Start™ Bradford Protein Assay Kit (#5000202, BIORAD) according to the manufacturer’s instruction^34^. Denaturing electrophoresis and western blotting were performed using the NuPAGE^®^ electrophoresis system (Invitrogen) following the manufacturer’s protocol. For all other Western blots, cells were lysed in lysis buffer (50mM Tris-HCl pH 7.8 (Sigma Aldrich), 100mM NaCl, 50mM KCl and 0.5% Triton X-100 (Sigma Aldrich), supplemented with cOmplete™ EDTA free Protease inhibitor cocktail tablet (Roche)) and prepared in Laemmli 2× buffer (Sigma-Aldrich). Cell proteins were resolved by SDS-PAGE using Mini-PROTEAN TGX Precast Gels (Bio-Rad). Immunoblotting was carried out using the following primary antibodies: rabbit α-ANP32A (Sigma-Aldrich #AV40203), mouse α-β-actin (Sigma-Aldrich #A2228), mouse α-FLAG (F1804, Sigma-Aldrich), mouse α-Lamin B1 (MAB5492, Merck), mouse α-PCNA (c907, Santa Cruz), rabbit α-Histone 3 (AB1791, Abcam), rabbit α-vinculin (AB129002, Abcam), rabbit α-*Gaussia* Luc (E80235, NEB), rabbit α-PB1 (PA5-34914, Invitrogen) and rabbit α-PB2 (GTX125926, GeneTex). The following secondary antibodies were used: goat anti-rabbit HRP (CST #7074), antimouse HRP (CST #7076), goat α-mouse AlexaFluor-568 (A11031, Invitrogen), sheep α-rabbit HRP (AP510P, Merck) and goat α-mouse HRP (STAR117P, AbD Serotec). Protein bands were visualized by chemiluminescence (ECL+ western blotting substrate, Pierce) using a FUSION-FX imaging system (Vilber Lourmat).

### Quantification of chANP32A, B and E mRNA levels

Total RNA from PGC fibroblast and DF-1 cells were extracted using an RNeasy mini kit (Qiagen), following manufacturer’s instructions. During extraction of RNA, RNeasy columns were treated with RNase-Free DNase (Qiagen). RNA samples were quantified using a Nanodrop Spectrophotometer (Thermo Scientific). Equal concentrations of RNA were subject to first strand synthesis using RevertAid (Thermo Scientific) with Oligo(dT) (Thermo Scientific). This product was then quantified with Fast SYBR™ Green Master Mix (Thermo Scientific) using the following sequence-specific primer pairs: RS17, (5′-ACACCCGTCTGGGCAACGACT-3′ and 5′-CCCGCTGGATGCGCTTCATCA-3′), RPL30 (5′- CCAACAACTGTCCTGCTTT-3′ and 5′-GAGTCACCTGGGTCAATAA-3′), chANP32A (5′- GTTTGCAACTGAGGCTAAGC-3′ and 5′-CAACTGTAGGTCATACGAAGGC-3′), chANP32B (5′- GGTGGCCTTGAAGTTCTAGC-3′, and 5′-ATGAGCATCGTCACCTCGC-3′), chANP32E (5′- GAACTAGAGTTTCTTAGCATGG-3′ and 5′- TCTCTCTGCAAGGACCTCCAG-3′). Real-time quantitative PCR analysis was performed (Applied Biosystems ViiA 7 Real-Time PCR System).

### Safety/biosecurity

All work with infectious agents was conducted in biosafety level 2 facilities, approved by the Health and Safety Executive of the UK and in accordance with local rules, at Imperial College London, UK.

### Bioinformatics

ANP32 sequences were downloaded from Ensembl (Gene Trees ENSGT00940000153254 and ENSGT00940000154305.) Amino acid sequences were aligned using MUSCLE^35^ and the maximum likelihood tree was constructed using RAxML-HPC2 v.8.2.10^36^ (GTRGAMMA model, 100 bootstraps) on XSEDE run on CIPRES^37^. Mapmodulin from *Drosophila melanogaster* was used as an outgroup.

## Supplementary Information

Supplementary Figure 1. Phylogenetic and sequence analysis reveals avian ANP32B to be a paralog of mammalian ANP32B.

Full tree used to make the cladogram in Figure 1. This tree was made using RAxML with 100 bootstraps and mapmodulin as an outgroup.

Supplementary Figure 2. Synteny of ANP32 genes.

Chromosome locations of ANP32 family members A, B, C and E from coelacanth (*Latimeria chalumnae*, Xenopus (*Xenopus tropicalis*), chicken (*Gallus gallus*), zebra finch (*Taeniopygia guttata*), opossum (*Monodelphis domestica*), mouse (*Mus musculus*) and Human (*Homo sapiens*).

Supplementary Figure 3. Sequence analysis of ANP32 in genome edited chicken cells.

**a.** DNA sequence analysis of ANP32B from genomic DNA of DF-1 WT and ΔB clones, showing the target sequence of the gRNA pair used in the CRISPR/Cas9 reaction. Allele A had a 16bp deletion and allele B a 40bp deletion, resulting in premature stop codons in the ANP32B sequence. Alignment of DNA sequence from WT, Δ33 and KO PGCs, showing the target sequence of the gRNAs used in the CRISPR/Cas9 reaction. **b.** Comparison between WT and KO PGCs showed an 8bp deletion in exon 1 of ANP32A in KO PGC cells, resulting in a truncated ANP32A protein. **c.** Intron and exon 5 sequence comparison of WT and Δ33 cells revealed a 400bp deletion resulting in the loss of exon 5. **d.** qRT-PCR analysis of mRNA isolated from WT and bKO DF-1 cells. Data are Δct of RPL30, ANP32A, B or E to RS17. **e.** qRT-PCR analysis of mRNA isolated from WT, Δ33 or aKO PGC derived fibroblast cells. Data are Δct of RS17, ANP32A, B or E to RPL30. Annotated alignments generated using Geneious R6 software.

Supplementary Figure 4. *In vitro* reprogramming of chicken PGCs into adherent fibroblast-like cells.

a. Diagram describing the method to differentiate PGCs from day 3 chicken embryos. b. Fluorescent images of live cells demonstrating the differentiation of GFP^+^ PGCs into fibroblast-like cells.

Supplementary Figure 5. Nuclear localisation of exogenously expressed ANP32 proteins in 293T cells.

Immunofluorescent images of 293T cells expressing ANP32 constructs fixed and stained with DAPI to highlight the nucleus. a. FLAG-tagged ANP32A constructs were imaged by probing with mouse α-FLAG antibody and detected by α-mouse AlexaFluor-568: Empty vector(1), huANP32B(2), huANP32B^33^(3), chANP32B(4), chANP32B^33^(5), huANP32B^33^_LRR_(6), huANP32B^33^_CENT_(7), huANP32B^33^_LCAR_(8), huANP32B^33^_N+LRRi_(9), huANP32B^33^_LRR2+3_(10), huANP32B^33^_LRR4+s_(11), chANP32A(12), chANP32A_scr149-175_(13), chANP32A_scr176-208_(14). b. ANP32A with mCherry fused to the C-terminus: Empty Vector(1), chANP32A(2), chANP32B(3), chANP32A_K116H_(4), chANP32A_N127M_(5), chANP32A_N129I_(6), chANP32A_D130N_(7), chANP32A_K137T_(8). All images were prepared using ImageJ^26^ and Microsoft PowerPoint.

Supplementary Figure 6. Avian ANP32B proteins share the I129 and N130 residues in LRR5.

Alignment comparing sequence of LRR5 ANP32B sequence from *Homo sapiens* and 22 avian species (residues 115 to 141). Protein sequences downloaded from NCBI and aligned using Geneious R6 software.

Supplementary Figure 7. Western blot analysis of split luciferase constructs.

**a.** Normalised luciferase Ratio was calculated by the equation described in this diagram, whereby the bioluminescence measured by interacting partners A and B fused to luc1 and luc2 are divided by the sum of the bioluminescence of the unfused controls. **b.** Western blot analysis of luc-fused constructs (α-Vinculin, PB1, *Gaussia* luciferase and histone 3)

