## Supplementary Information for "Avian ANP32B does not support influenza A virus polymerase and influenza A virus relies exclusively on ANP32A in chicken cells"

**Supplementary Figure 7. Western blot analysis of split luciferase constructs.**

**a.** Normalised luciferase Ratio was calculated by the equation described in this diagram, whereby the bioluminescence measured by interacting partners A and B fused to luc1 and luc2 are divided by the sum of the bioluminescence of the unfused controls. **b.** Western blot analysis of luc-fused constructs (α-Vinculin, PB1, *Gaussia* luciferase and histone 3).

Supplementary Figure 1

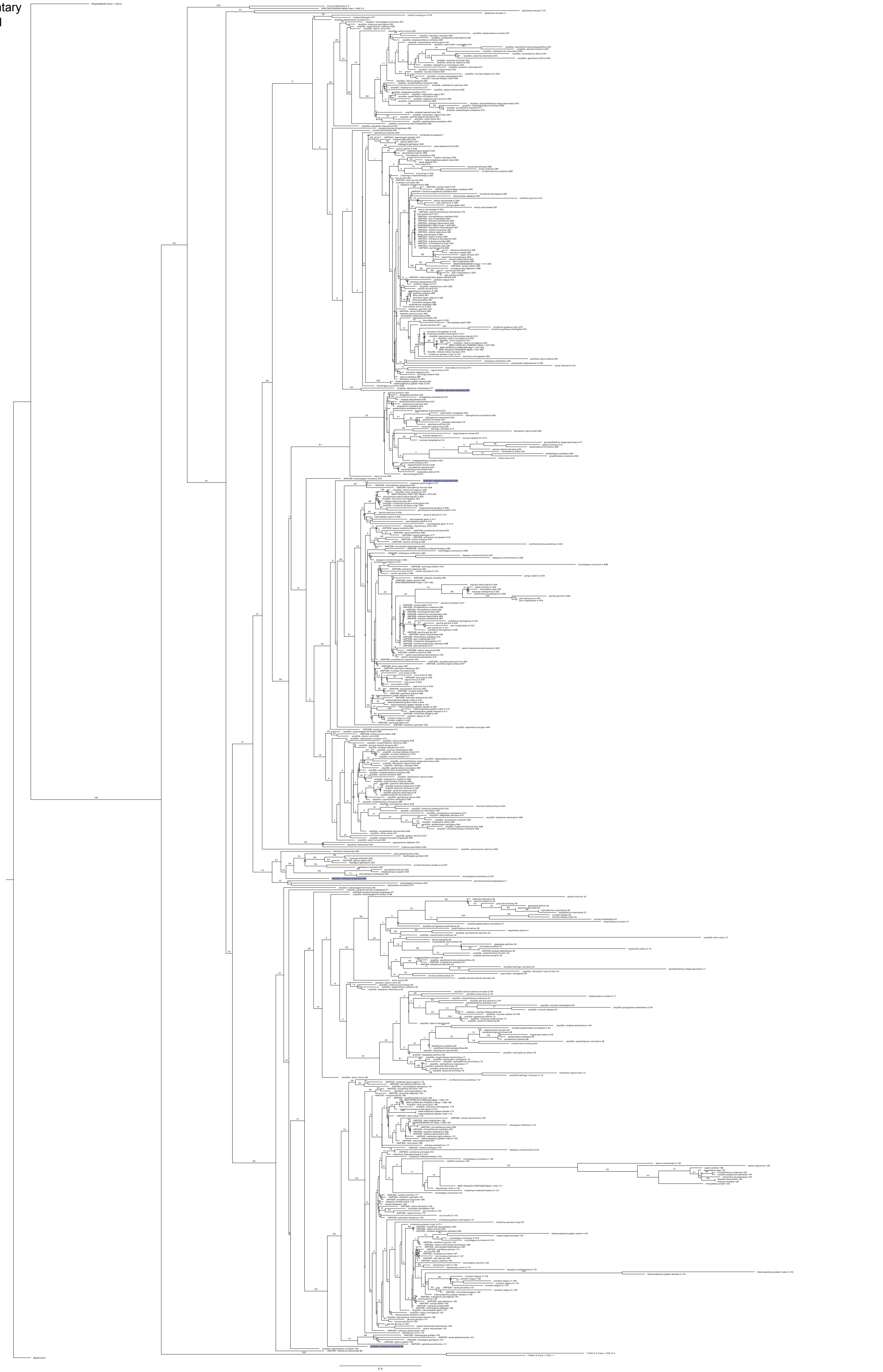

Supplementary Figure 2

***ANP32A***

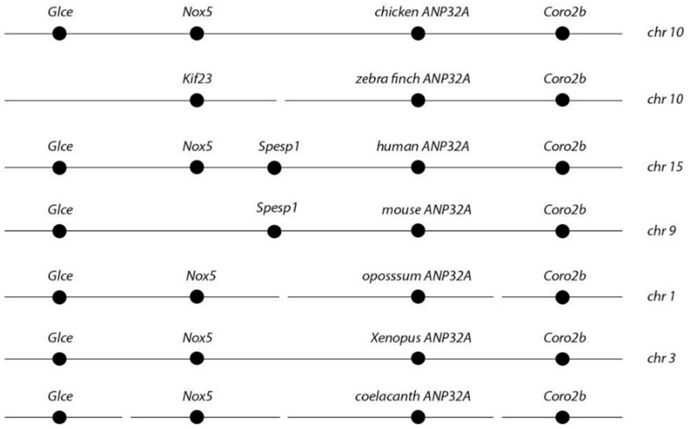


***ANP32B***

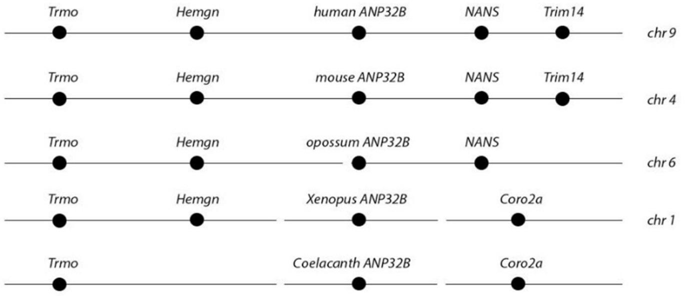


***ANP32C***

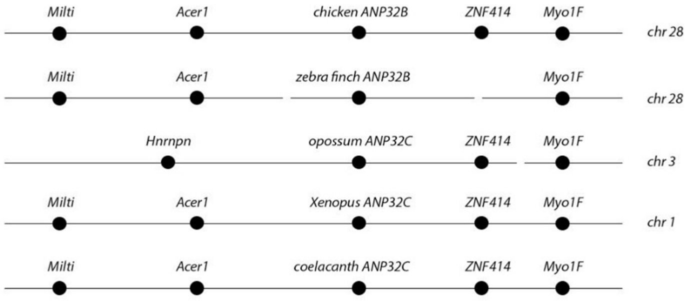


***ANP32E***

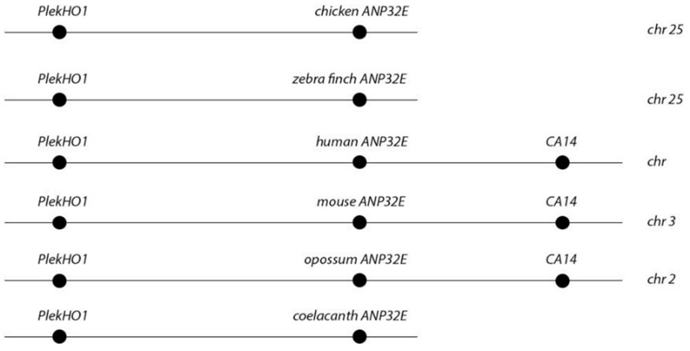

Supplementary Figure 3

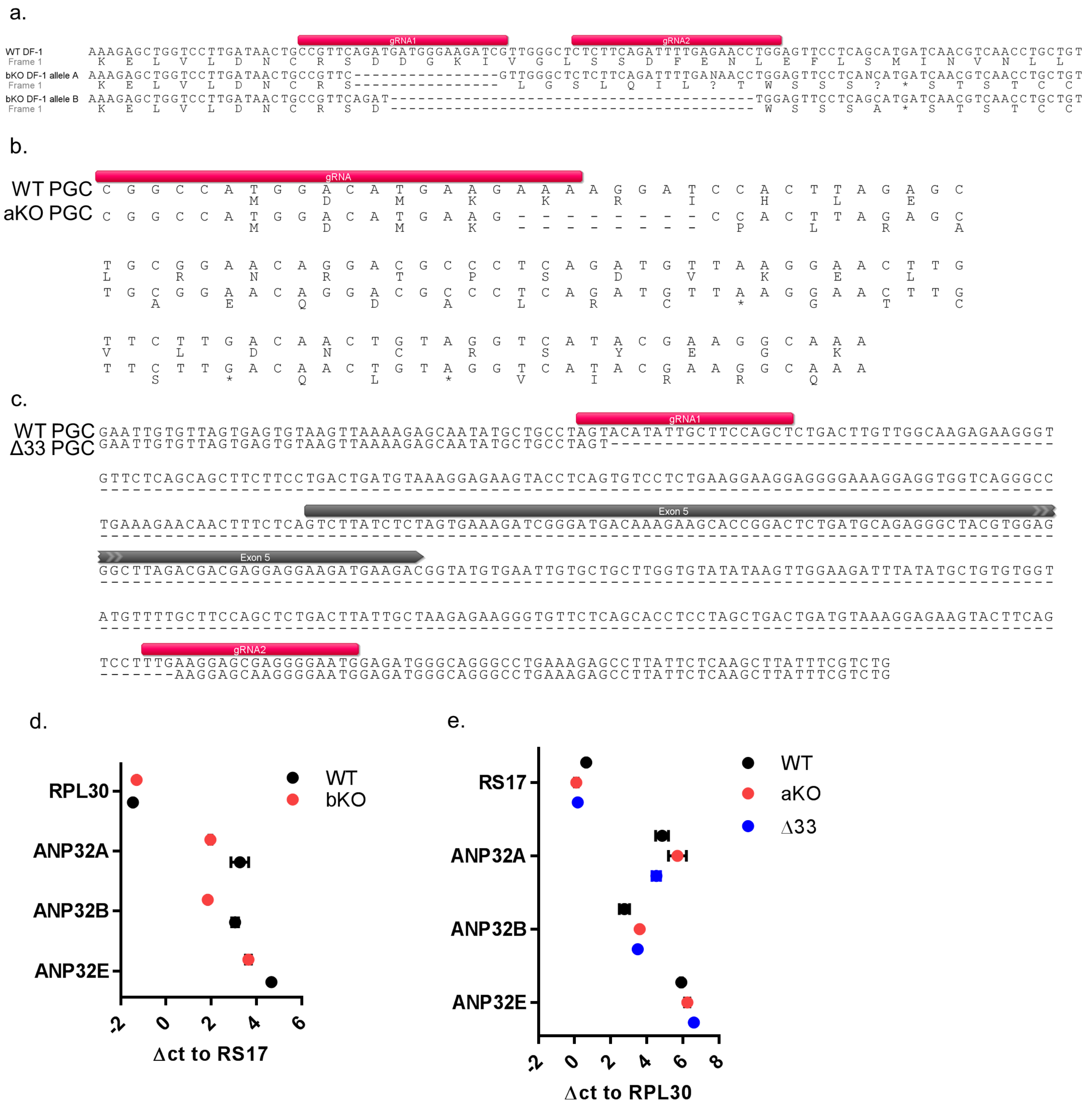

Supplementary Figure 4

a.

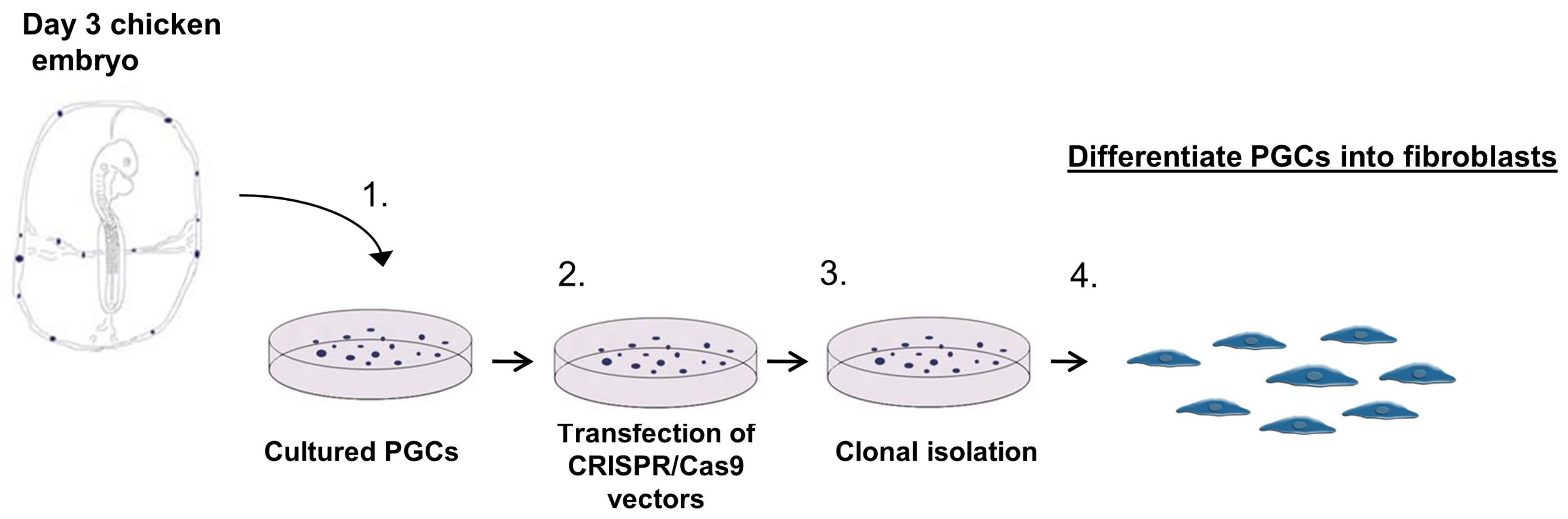


b.

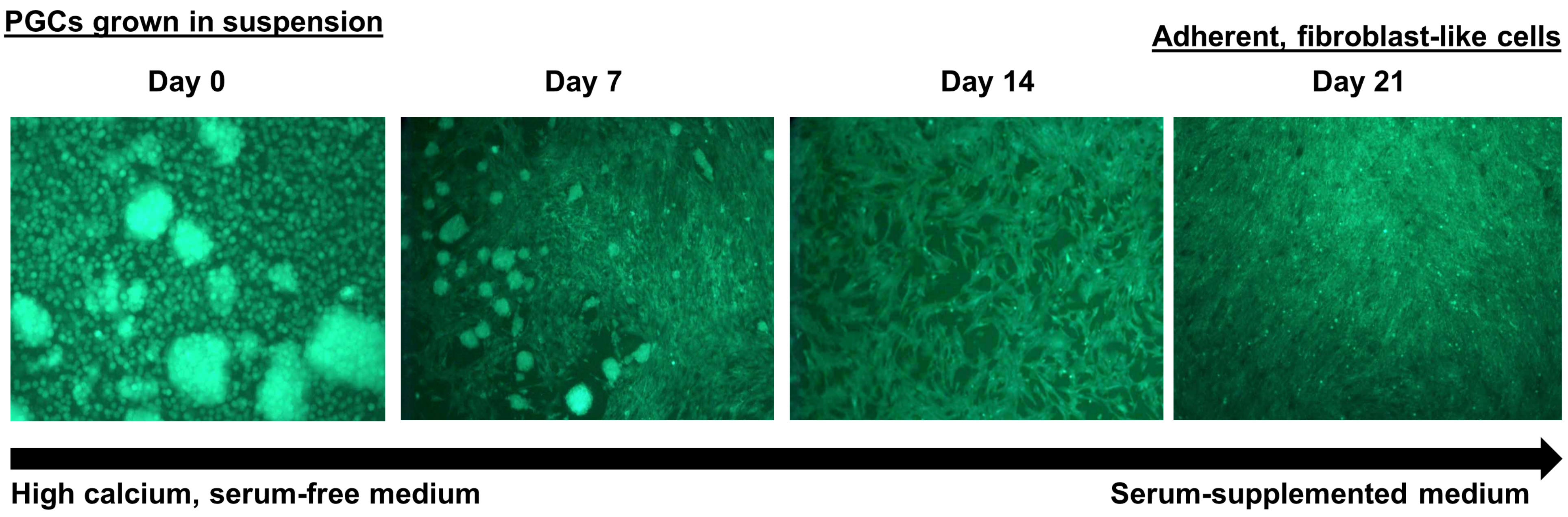

Supplementary Figure 5

a

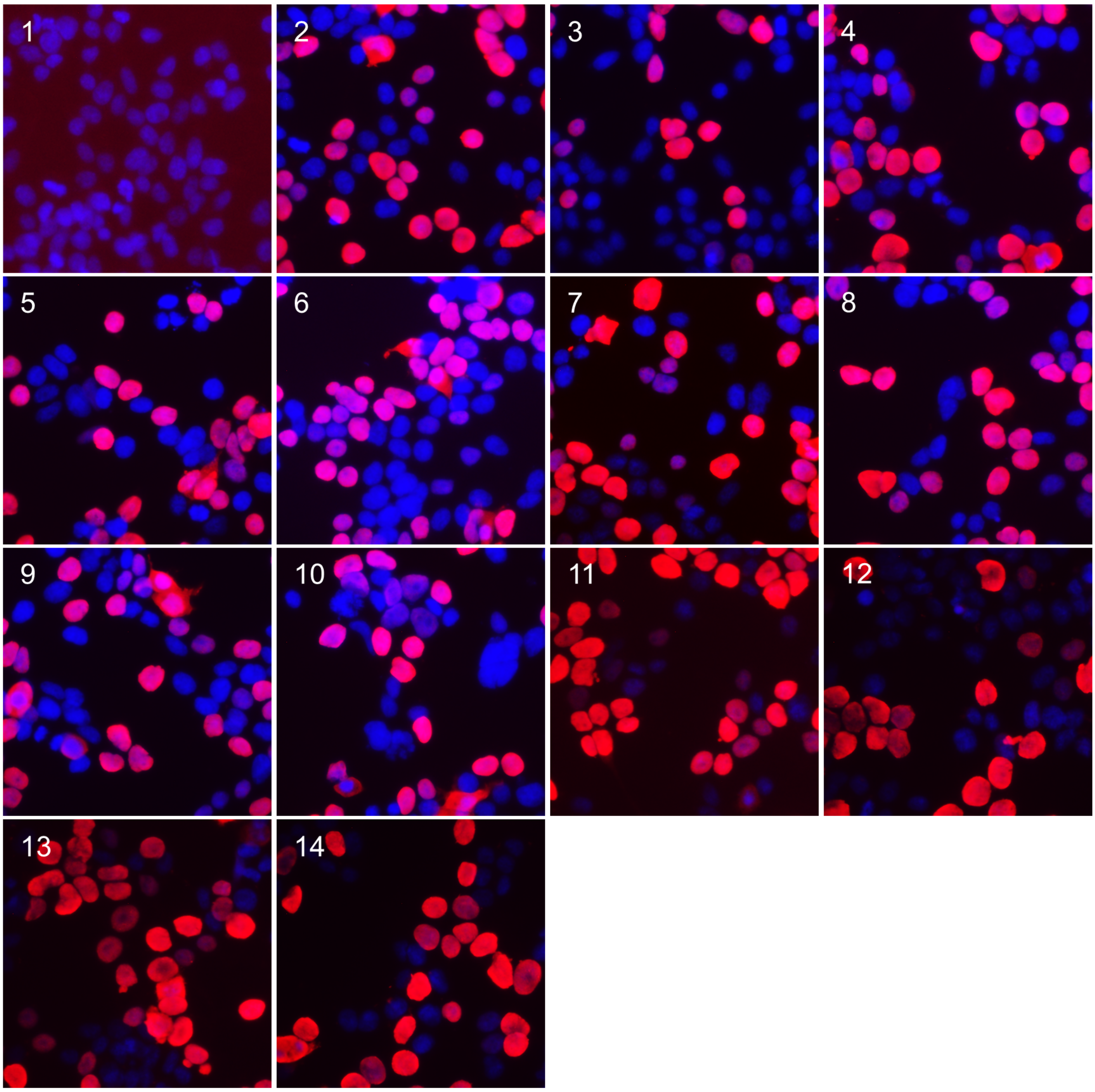


b

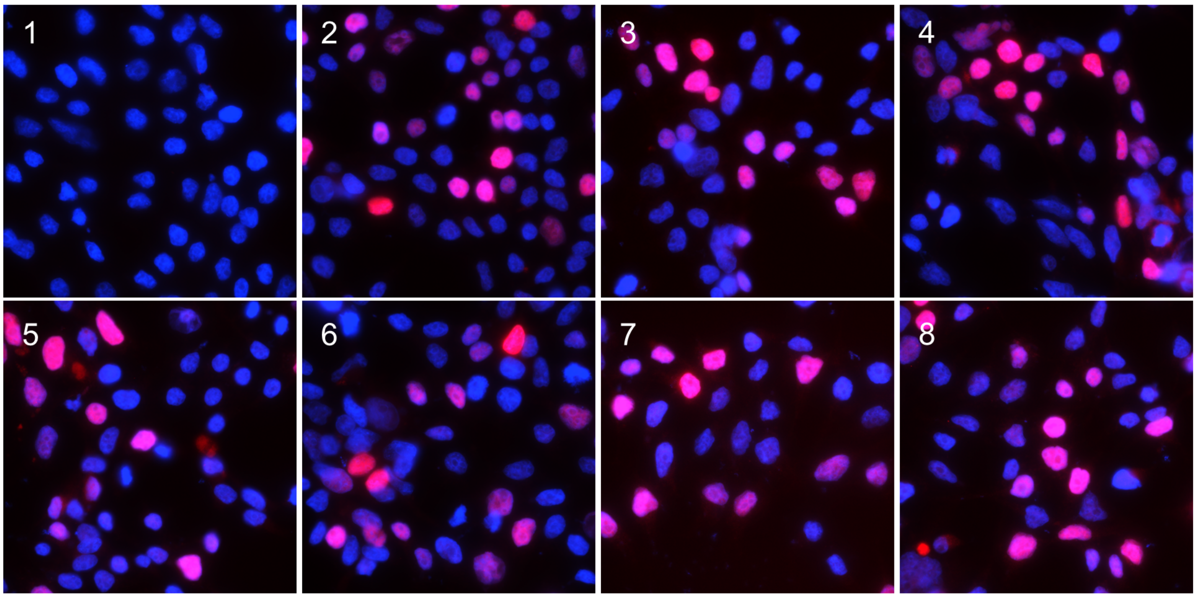

Supplementary Figure 6

```
                                                     116         127 129 130      137
                                                      |           |   |  |         |
Consensus                                            LHSLDLFNCEVTMLINYRESVFTLLPQ
Homo sapien [Q92688.1]                               .K..........N.ND......K....
Gallus gallus [XP_015155299.1]                       ...........................
Acanthisitta chloris [XP_009074950.1]                ......................A....
Anas platyrhynchos [XP_012963723.1]                  ...........................
Apteryx australis mantelli [XP_013806136.1]          ...........................
Aquila chrysaetos canadensis [XP_011594284.1]        ...........................
Calidris pugnax [XP_014817120.1]                     ...........................
Chlamydotis macqueenii [XP_010127925.1]              ...........................
Coturnix japonica [XP_015742181.1]                   ...........................
Cyanistes caeruleus [XP_023798892.1]                 ......................A....
Ficedula albicollis [XP_016160035.1]                 ......................A....
Haliaeetus albicilla [XP_009914439.1]                ...........................
Haliaeetus leucocephalus [XP_010560591.1]            ...........................
Lepidothrix coronata [XP_017691899.1]                ......................A....
Lonchura striata domestica [XP_021399982.1]          .R....................A....
Meleagris gallopavo [XP_010723174.1]                 ...........................
Melopsittacus undulatus [XP_012986151.1]             ...........................
Nestor notabilis [XP_010013914.1]                    ...........................
Numida meleagris [XP_021234318.1]                    ...........................
Pseudopodoces humilis [XP_005531306.1]               ......................A....
Sturnus vulgaris [XP_014738541.1]                    ......................A....
Tinamus guttatus [XP_010213231.1]                    .R.........................
Tyto alba [XP_009973260.1]                           ...........................
```

Supplementary Figure 7

a

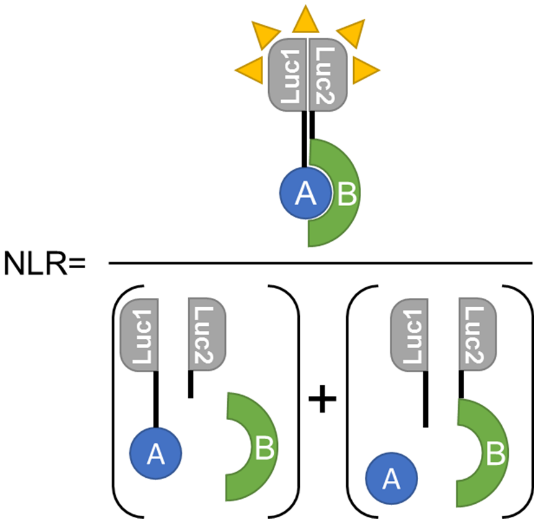


b

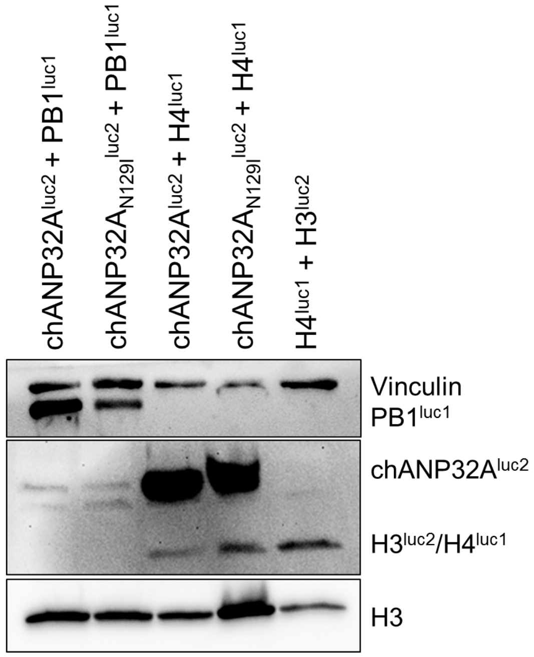
